# Survival Genie, a web platform for survival analysis across pediatric and adult cancers

**DOI:** 10.1101/2021.09.28.462224

**Authors:** Bhakti Dwivedi, Hope Mumme, Sarthak Satpathy, Swati S. Bhasin, Manoj Bhasin

**Affiliations:** Winship Cancer Institute, Emory University, Atlanta, GA; Aflac Cancer and Blood Disorders Center, Children’s Healthcare of Atlanta, Atlanta, GA; Department of Pediatrics, Emory University, Atlanta, GA; Department of Biomedical Informatics, Emory University, Atlanta, GA

**Author notes:** **Corresponding author**, Manoj K. Bhasin, MS, Ph.D. Aflac Cancer and Blood Disorders Center, Children Healthcare of Atlanta, Woodruff Memorial Research Building, Room 4107, 101 Woodruff Circle, 4th Floor East, Emory School of Medicine, Atlanta, GA 30322.

**Keywords:** survival analysis, gene set, single-cell RNA sequencing, cancer

## Abstract

The genomics data-driven identification of gene signatures and pathways has been routinely explored for predicting cancer survival and making decisions related to targeted treatments. A large number of packages and tools have been developed to correlate gene expression/mutations to the clinical outcome but lack the ability to perform such analysis based on pathways, gene sets, and gene ratios. Furthermore, in this single-cell omics era, the cluster markers from cancer single-cell transcriptomics studies remain an underutilized prognostic option. Additionally, no bioinformatics online tool evaluates the associations between the enrichment of canonical cell types and survival across cancers. Here we have developed Survival Genie, a web tool to perform survival analysis on single-cell RNA-seq (scRNA-Seq) data and a variety of other molecular inputs such as gene sets, genes ratio, tumor infiltrating immune cells proportion, gene expression profile scores, and tumor mutation burden. For a comprehensive analysis, Survival Genie contains 53 datasets of 27 distinct malignancies from 11 different cancer programs related to adult and pediatric cancers. Users can upload scRNA-Seq data or gene sets and select a gene expression partitioning method (i.e., mean, median, quartile, cutp) to determine the effect of expression levels on survival outcomes. The tool provides comprehensive results including box plots of low and high-risk groups, Kaplan-Meier plots with univariate Cox proportional hazards model, and correlation of immune cell enrichment and molecular profile.

The analytical options and comprehensive collection of cancer datasets make Survival Genie a unique resource to correlate gene sets, pathways, cellular enrichment, and single-cell signatures to clinical outcomes to assist in developing next-generation prognostic and therapeutic biomarkers. Survival Genie is open-source and available online at https://bbisr.shinyapps.winship.emory.edu/SurvivalGenie/.

## INTRODUCTION

Over the past decade advancement in genomics and epigenomics technologies has produced a large amount of data to understand the complex mechanisms of cancer. This accessibility and affordability of genome sequencing resulted in the completion of multiple ambitious projects in the cancer arena such as The Cancer Genome Atlas (TCGA)^1^, International Cancer Genome Consortium (ICGC)^2^, and The Therapeutically Applicable Research to Generate Effective Treatments (TARGET)^3^ to generate a comprehensive molecular landscape of different cancers. The TCGA program has profiled genetic and epigenetics landscapes of ~20,000 tumor samples across 33 major adult cancer types to generate novel insights for better diagnosis and treatment of cancer. Similarly, the TARGET^3^ program has generated comprehensive genetics, epigenetics, and transcriptome profiles of 6,000 samples from nine pediatrics tumor types to understand molecular mechanisms and develop novel therapies. These ambitious projects provided access to omics, clinical, demographic, and outcome information through Genomic Data Commons (GDC) gateway^4^ to understand the impact of various associated factors on cancer outcomes besides collecting molecular information. The correlative analysis of these vast *omics* and clinical data can assist in identifying robust prognostics biomarkers to predict cancer outcomes. However, given the inherent complexity of data coupled with the sheer volume from numerous initiatives, the overall potential is largely reduced due to the lack of systematic and user-friendly online analytical tools. Utilization by a larger cross-section of users of such a massive amount of data will require access to the data without the need to download the files and significant knowledge of analytics as well as programming languages such as python and R. This raises the need for the development of user-friendly online platforms for the exploration of this data.

To address this unmet need, multiple online platforms for exploring the association of genes and proteins with the clinical outcome have been developed^5^. Among the most used survival analysis tools are the cBioPortal^6^ and GDC data portal^4^, providing users exploratory analyses of multi-omics cancer datasets and survival analysis of single genebased on DNA alterations only. UALCAN^7^ is an interactive web resource for analyzing cancer OMICS data from TCGA, MET500, and CPTAC to correlate gene, protein, and miRNA expression profile and patient survival information. TRGAted^8^ tool allows survival analysis of single and multiple proteins across TCGA cancer types. This tool has clinical options with multiple optimal cut-offs; however, is limited to protein data and uses the mean expression as a cut-point for querying multiple proteins. Likewise, the KMplotter ^9,10^ allows studying the clinical impact of individual genes in cancers from TCGA, GEO, and EGA using different splitting methods (e.g., median, percentile, quartiles). In this tool, users can upload multiple genes, ratios of two genes, and mean or median expression of genes, yet analysis will be performed at the individual gene level. The tool also allows restrictive analysis based on clinical subtypes and the proportion of immune cells. In this series, LOGpc^11^ is another web server that estimates survival in 27 cancer types from TCGA and GEO using RNA-seq data but limited to single gene-based analysis. A recently developed tool, ESurv^12^ also provides univariate survival analysis by a single gene or cancer type from TCGA datasets.

Multiple genes, gene sets, and canonical pathways play a key role in defining the different subtypes of cancers with varying aggressiveness and outcomes. This raises the need for a tool to perform survival analysis from a set of genes rather than on a single gene basis within a set. Keeping this in mind, we developed the survival Genie platform by compiling a comprehensive list of adult and pediatric cancers along with the implementation of multiple approaches for prognostic genomics features identification. For comprehensive survival evaluation, the Survival Genie contains 53 datasets of 27 distinct malignancies from 11 different cancer programs for both adult and pediatric cancers. Users can upload single-cell data or gene sets and select partitioning methods (i.e., mean, median, quartile, and cutp) to determine the effect of expression levels on survival outcomes. The tool provides comprehensive results including box plots of low and high-risk groups, Kaplan-Meier plots with univariate Cox proportional hazards model, and correlation of immune cell enrichment and molecular profile.

## METHODS

### Collection and processing of cancer data

Survival Genie contains 53 datasets of 27 distinct malignancies from 11 different cancer programs for both adult and pediatric cancers (Supplementary Table 1). All the clinical and genomics (transcriptomic and mutation) data were downloaded from the GDC data portal^4^ using The GenomicDataCommons Bioconductor R package^13^. The samples with paired survival (e.g., days to follow-up, days to death, vital status), mRNA-seq, and Whole Exome Sequencing are only used for survival analysis. The three tumor types (primary, recurrent, and metastatic) are included in the analysis depending on the cancer type. Additionally, for correlating the survival information with immune infiltrate, the clinical Histology and Eosin (H&E) digital image-based quantification of Tumor-infiltrating lymphocytes (TILs) were obtained from the Cancer Imaging Archive^14^ for the 13 TCGA datasets.

### Survival analysis

Survival Genie performs Overall survival (OS) and Event-free survival (EFS) statistical analysis as implemented in the ‘survival’ R package ^15^. Kaplan Meier survival curves are used to estimate the OS/EFS using the survfit function and a log-rank test is done to compute differences in OS/EFS between the defined high- and low-risk groups. Univariate analysis with Cox proportional hazards regression model is performed on the patient data using coxph function in R/Bioconductor ^16^.The survival association is considered significant if the p-values for the log-rank and Wald tests are lower than 0.05.

### Implementation

The Survival Genie source code is written in the R programming language ^17^ and the interactive web-server is implemented using the shiny R package ^18^. Cancer clinical and genomics datasets for survival analyses are processed and analyzed using Perl and R languages. The tool has been extensively tested on multiple operating systems (Linux, Mac, Windows) and web-browsers (Chrome, Firefox, and Safari). The tool is currently hosted on a 64bit CentOS 6 backend server running the Shiny Server program designed to host R Shiny applications ^18^. The tool along with source code is also available on the GitHub repository https://github.com/bhasin-lab/SurvivalGenie. The detail step-wise analysis tutorial about the tool is also available on YouTube at https://www.youtube.com/watch?v=H5s6OYvwwoo.

## RESULTS

### Development of Survival Genie

**Figure 1** shows the overview of the user-friendly Survival Genie web interface for choosing analysis type, inputs, cancer datasets, parameters to explore and visualize the effect of dichotomous molecular (e.g., Gene) profiles on patient survival outcomes. Detailed documentation for input and output parameters, and example data are available on the help page of Survival Genie web server.

**Figure 1:**
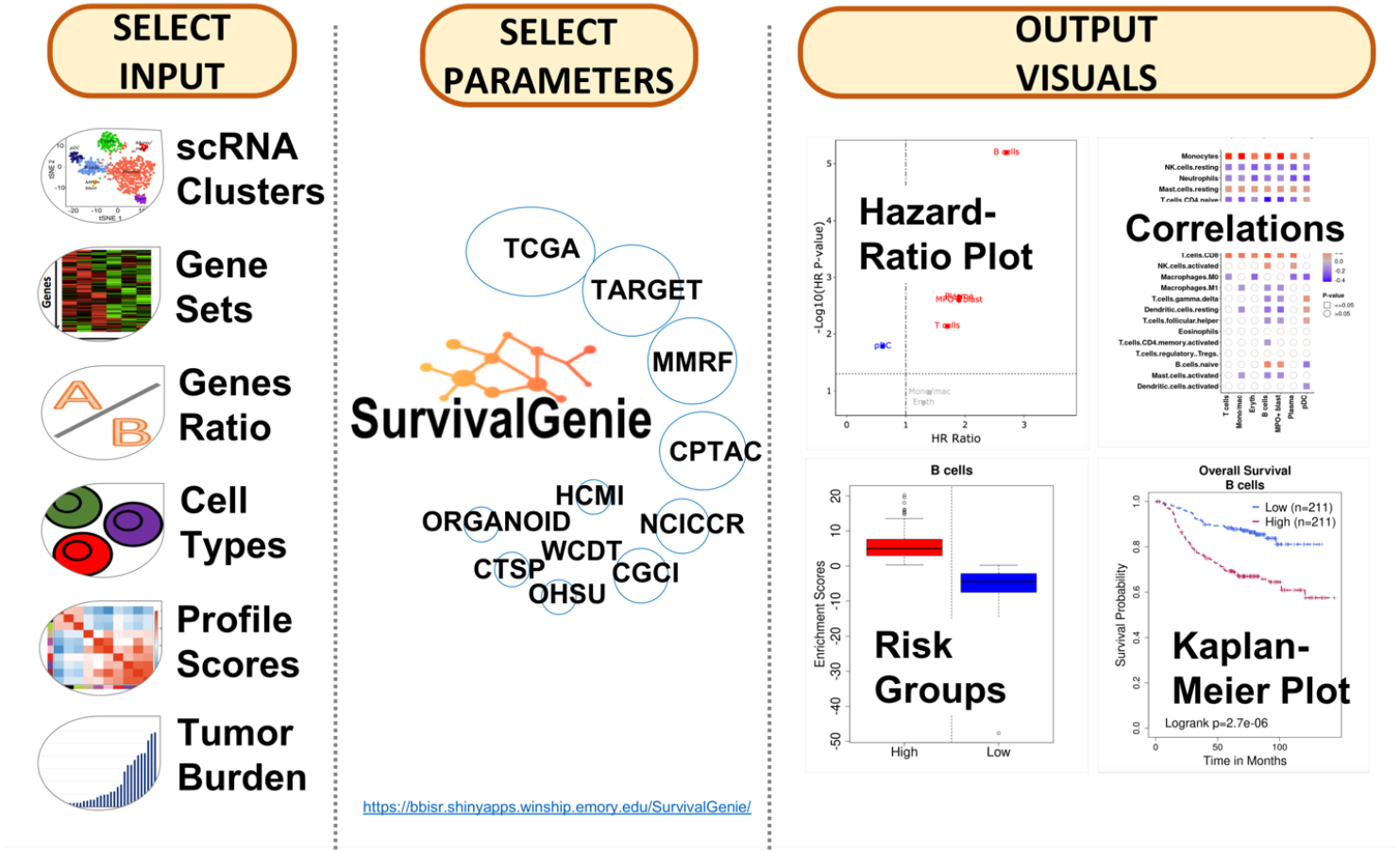
An overview of Survival Genie platform showing input, analysis parameters, and output.

#### Pathways, Gene sets, and Systems Biology modules-based survival analysis

Survival Genie provides a range of input options (i.e., gene-based, cell-based, profile-based, and mutation-based) to allow flexible and comprehensive survival analysis based on the user’s needs (**Fig. 2, steps 1-2**). **(a) Gene-based** input option requires HGNC approved gene symbols as input, the most popular and widely used option for identifying prognostic gene markers. ***Gene Sets*** allow the input of a list of genes from pathways or biological processes as a gene set. An aggregated enrichment score is computed for a gene set using the single-sample gene set enrichment method (ssGSEA ^19^) to estimate optimal cut-off in predicting survival outcomes. The gene set option also allows submission of **Single*-cell RNA-seq clusters markers*** file containing lists of significant gene markers for each identified cell cluster derived from any single-cell RNA-seq experiment using analysis tools such as Seurat ^20^. An adapted ssGSEA is implemented to compute enrichment scores (ES) for marker genes for identified clusters from singlecell experiment. Each tumor cluster enrichment score is then divided into two-risk categories to predict patient outcomes and of associations to tumor cell composition. *Additionally, **Genes Ratio*** input option allows users to perform survival analysis on the ratio of two-genes computed from the normalized expression values, and lastly ***(iii) Single Gene*** input option allows users to perform single gene queries to study associations with clinical outcomes. **(b) Cell-based** is to query cell types derived from ***(i)* CIBERSOFT TILs proportion** estimates of tumor-infiltrating lymphocytes (TILs) using LM6 and LM22 cell signature matrix ^21^. The proportions were estimated using bulk tumor normalized FPKM expression data for each tumor for cancer types following the CIBERSORT method^21^ and ***(ii)* Digital TILs percentage** retrieved based on H&E images from The Cancer Imaging Archive^14^. **(c) Profile-based** input option is to query **Weighted gene profiles** derived from weighted gene co-expression network analysis (e.g., 18-gene T-cell inflammation gene expression profile, the so-called Gene Expression Profile, GEP scores). The GEP score is calculated as the weighted sum of the normalized FPKM expression values of the gene signature for each tumor. The weightings for each HGNC approved gene symbol in the signature are provided by the user in the uploaded submission file. **(d) Mutation-based** input option is to query **Tumor Mutation Burden (TMB)** within a cancer dataset based on all, only non-synonymous or exonic somatic mutations per megabase (Mb) of the genome.

**Figure 2.**
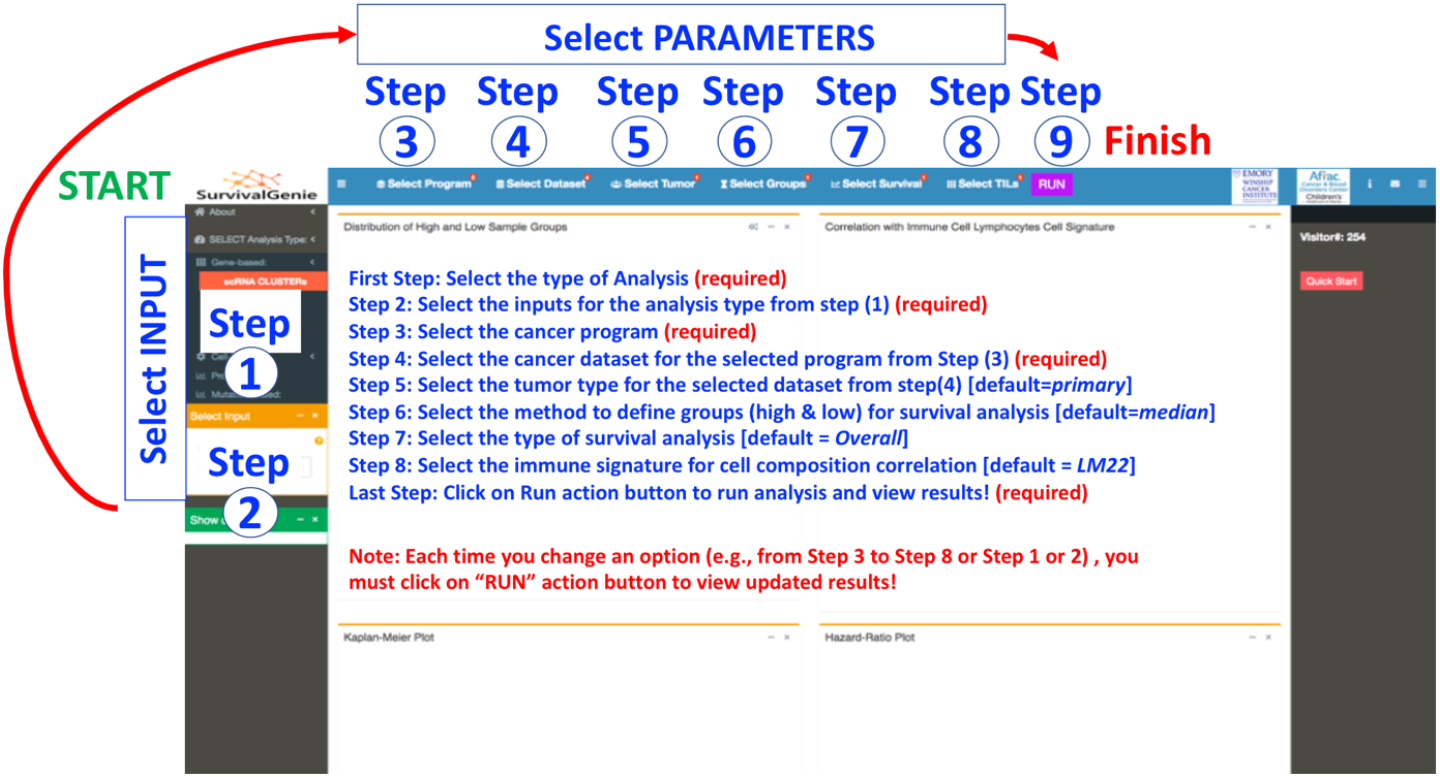
Summary of steps for performing survival analysis using Survival Genie Platform. The detailed instructions for analysis options and procedure are available in the demonstration tutorial video (https://www.youtube.com/watch?v=H5s6OYvwwoo) and in the user manual on the GitHub site (https://github.com/bhasin-lab/SurvivalGenie).

#### Cancer datasets, tumor types, optimal cut-offs, and survival options

In Survival Genie, users have options to select cancer program, tumor type, and methods for partitioning expression profile (**Steps 3-9, Fig. 2**). The comprehensive analysis parameter includes **(3) Selection of cancer program** from the available list that includes TCGA, TARGET, and MMRF CoMMpass. **(4) Selection of cancer datasets from selected cancer programs** to perform survival analysis. **(5) Selection of the tumor type,** primary, recurrent, or metastatic within the selected cancer dataset. The majority of cancer datasets consist of primary tumor samples with the exception of skin melanoma (TCGA-SKCM) that is dominated by metastatic tumor samples. **(6) Selection of partitioning methods** to define the dichotomous **patient** groups. There are four different partitioning methods (i.e., mean, median, percentile, and cutp) to **subset** the tumors into high and low groups to assess the association of expression levels with survival outcomes. *Percentile based* partitioning method allows the selection of desired upper and lower threshold (e.g., quartiles or 10^th^ vs. 90^th^ percentile) to subset tumor samples into low and high groups. The optimal cut-point *Cutp option* estimates the martingale residuals^6^ using the ‘survMisc’ ^22^ package to divide patients into high and low groups. **(7) Selection of survival analysis** from either overall or event-free survival time, although later is only applicable to pediatric cancer datasets from the TARGET program. **(8) Select TILs signature** based on 22 (LM22) or 6 (LM6) immune cell types. The relative fractions of cell types are estimated for LM6 and LM22 immune cell gene signature from bulk tumors FPKM gene expression data using the CIBERSORT deconvolution method ^21^. **(9) Submit analysis to perform analysis** after making appropriate selections of input parameters.

#### Analysis Outputs and Visualization

The results from the gene set and gene ratio based analysis are displayed in five separate tabs: ***(A)* Expression levels of high and low sample groups** displays either a box and whisker or bar plots showing the distribution of estimated molecular profile (e.g., gene expression, the expression ratio of two genes, gene set enrichment scores, cell proportion or mutation burden) **(Fig. 3A). (B) Correlation with lymphocytes cell signature** shows the colored grid matrix of Pearson correlation of CIBERSOFT deconvoluted lymphocytes RNA-seq gene expression data and the estimated molecular data profile (**Fig. 3B).** Correlation coefficients are indicated with a color gradient, blue color represents negative correlation, while red color represents the positive correlation. The shape denotes the significance of the correlation with squares showing significant associations (P-value <.05)**. (C) Kaplan-Meier (KM) plot** shows the KM survival curves in the stratified high (red) and low (blue) groups of patients with log-rank test (**Fig. 3C**). A log-rank p-value<0.05 is considered statistically significant. **(D) Hazard-Ratio (HR) plot** shows the survival significance by HR ratio and HR p-value (**Fig. 3D**). HR is estimated based on a univariate cox proportional hazard regression model with Wald-test. An HR value above 1 indicates an increase in hazard (poor outcome) while an HR value below 1 indicates a reduction in the hazard (good outcome), and an HR value equal to 1 indicates no effect. A p-value<0.05 is considered statistically significant. **(E) The Forest plot** shows the detailed table of the Univariate Cox-Regression survival analysis (**Fig. 3E**). The plot shows the hazard ratio and 95% confidence intervals associated with two groups considered in the univariable analysis along with Wald-test and log-rank p-values. The table also shows the cut-off values applied to subset the patients into high and low groups along with the sample numbers in each group. The squares represent the HR, and the horizontal lines depict the upper and lower limits of the HR 95% confidence interval. Significant associations are shown in red-filled HR value squares. An arrow at the end of the horizontal line indicates the higher upper limit of the 95% confidence interval than the maximum shown (i.e., 3). Additionally, the “**Show Output**” option (**Fig. 3**) allows users to select and view the plots (A) and (C) based on input clusters (for scRNA cluster analysis) or datasets (for single-gene analysis).

**Figure 3.**
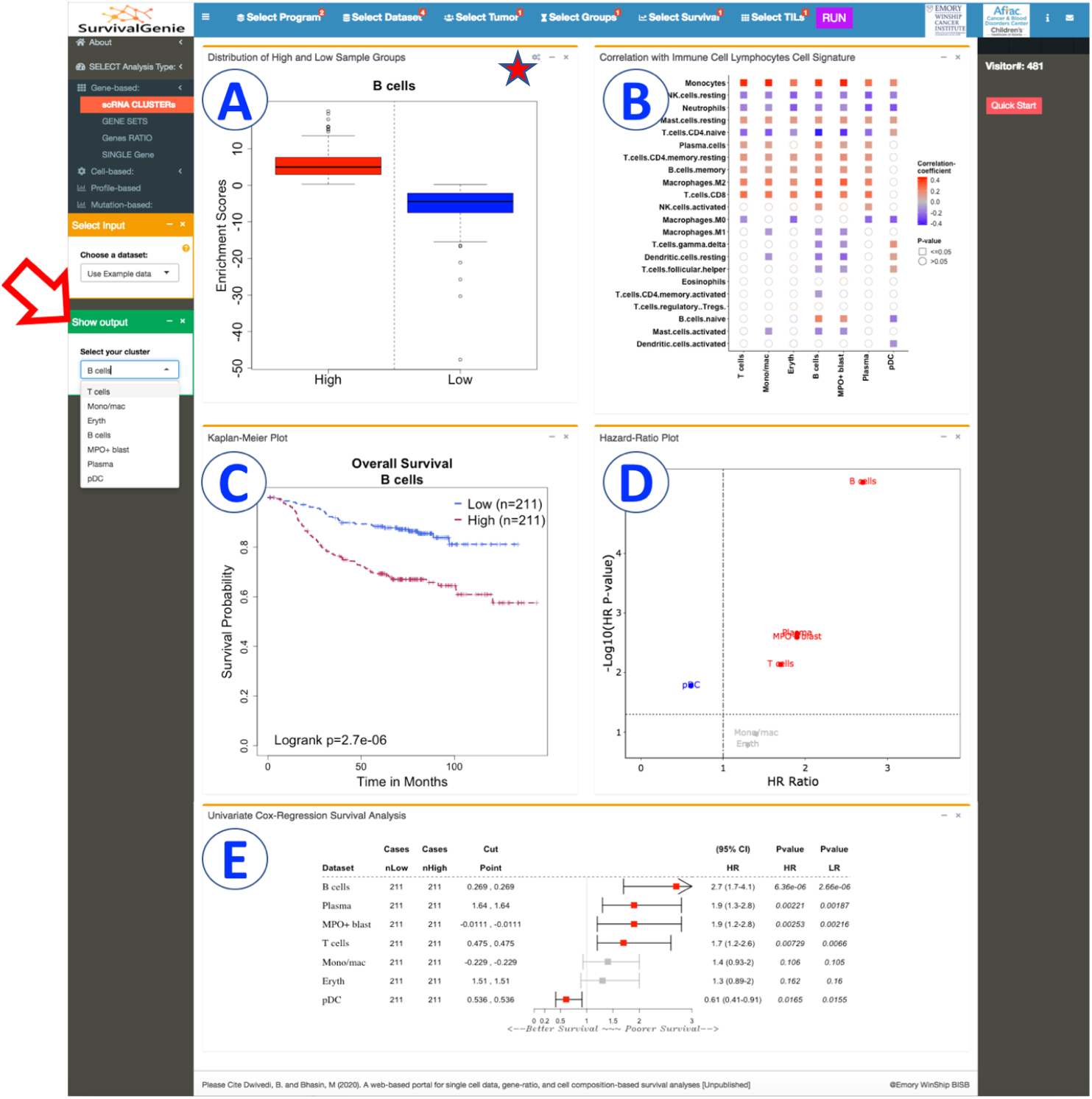
Summary of Survival Genie gene set analysis showing the five tabulated outputs. A. Distribution of Gene expression from high and low sample groups, B. Correlation with immune cell signature, C. Kaplan Meier, D. Hazard ratio, and E. Forest plot.

### Use Case Examples using Survival Genie

#### Use Case 1: Survival analysis of Hallmark gene sets

We performed Cox proportional hazards regression analysis to examine the correlation between hallmark gene sets (n=50) obtained from The Molecular Signatures Database (MSigDB)^23,24^ and survival (OS) across pediatric and adult cancers. Single sample enrichment scores were computed for each hallmark gene set across all cancer types using the ssGSEA method. The “survival” R package was utilized to calculate log-rank *P* values, hazard ratios (HR), and 95% confidence intervals (CI). Hierarchical clustering analysis (HCA) was performed on HR ratios to determine survival patterns of hallmarkgene sets (n=50) and cancer type (**Fig. 4,** Supplementary Table 2).

**Figure 4.**
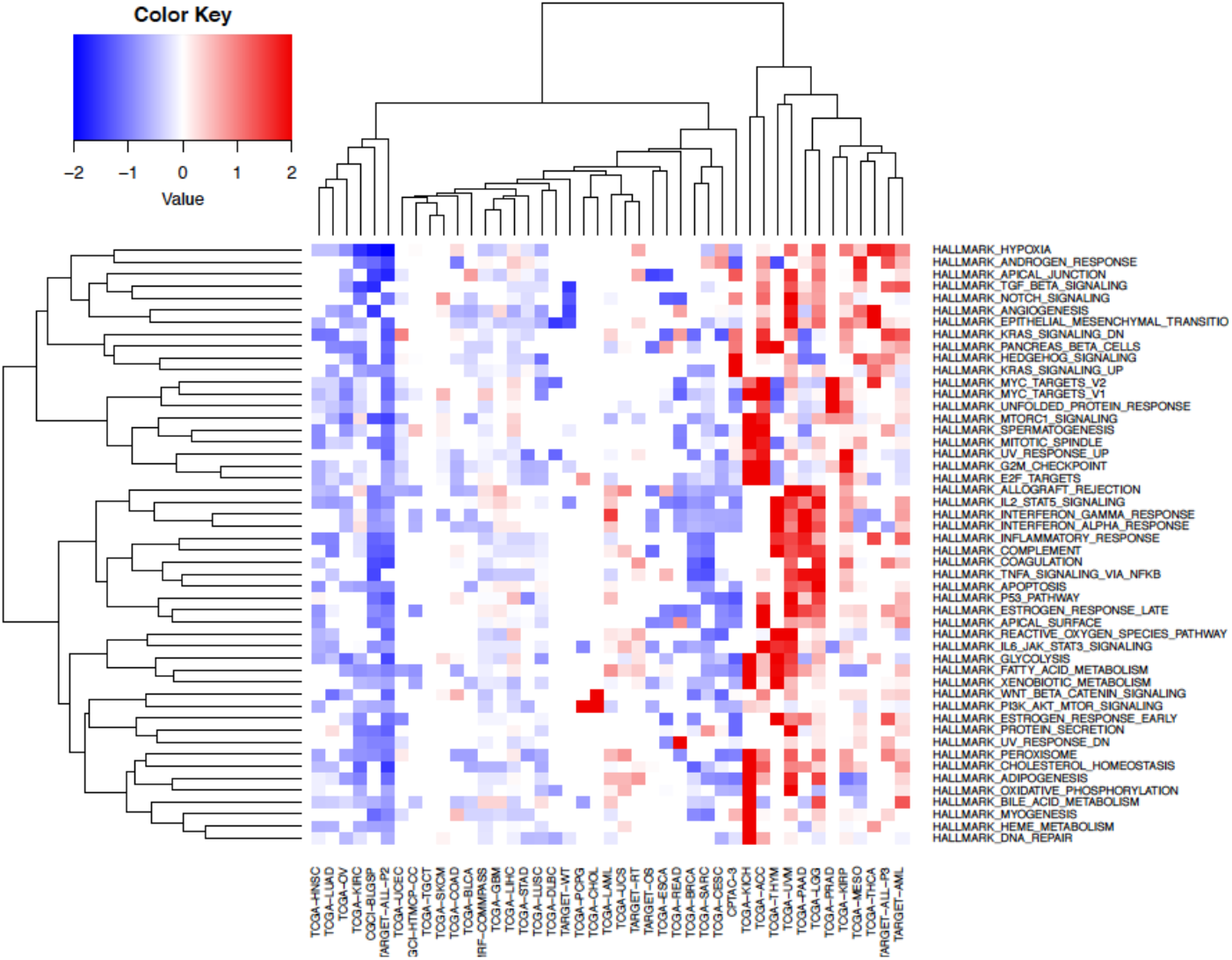
Gene sets based survival analysis. Hierarchical clustering of Broad hallmark-associated gene sets and cancer types. The predicted survival outcomes are visualized using Euclidean distance and complete linkage method, using Hazard Ratios values for each hallmark associated gene set (n=50) across all cancer datasets(n=43)

The HCA on hallmark gene sets HR ratio segregated cancers into three major groups (i.e., good, poor, and mixed outcomes) based on survival associations (**Fig. 4**). Interestingly across all the hallmark gene sets, pediatric lymphoblastic leukemia (i.e., TARGET-ALL-P2) shows a consistent pattern of overall good survival whereas TCGA low grade glioma (TCGA-LGG) showed overall bad survival. The association might be driven by overall low relapse rate for pediatric ALL and bad/short survival associated with low-grade glioma. The chronic inflammatory response-related gene sets such as “IL2 STAT5 signaling”, “Interferon-gamma response”, “interferon-alpha response”, “inflammatory response”, “apoptosis”, and “p53 pathway” were found to be significantly associated with poor overall survival in across multiple adult cancers including TCGA-LGG, TCGA-UVM, and TCGA-PADD datasets.

#### Use Case 2: *Survival Association of genes from Hallmark gene sets*

We performed single-gene analysis to explore further whether gene set analysis outcomes are influenced at the individual gene level survival associations. Cox regression analysis was performed using the RNA-seq expression of genes (n=4383) from 50 hallmark gene sets. The results of survival analysis across 47 types of cancer for individual gene from hallmark gene sets listed in Supplemental Table 3. We computed the proportion of significant genes associated with patient survival from each hallmark gene set across tumor types (**Fig. 5**).

**Figure 5.**
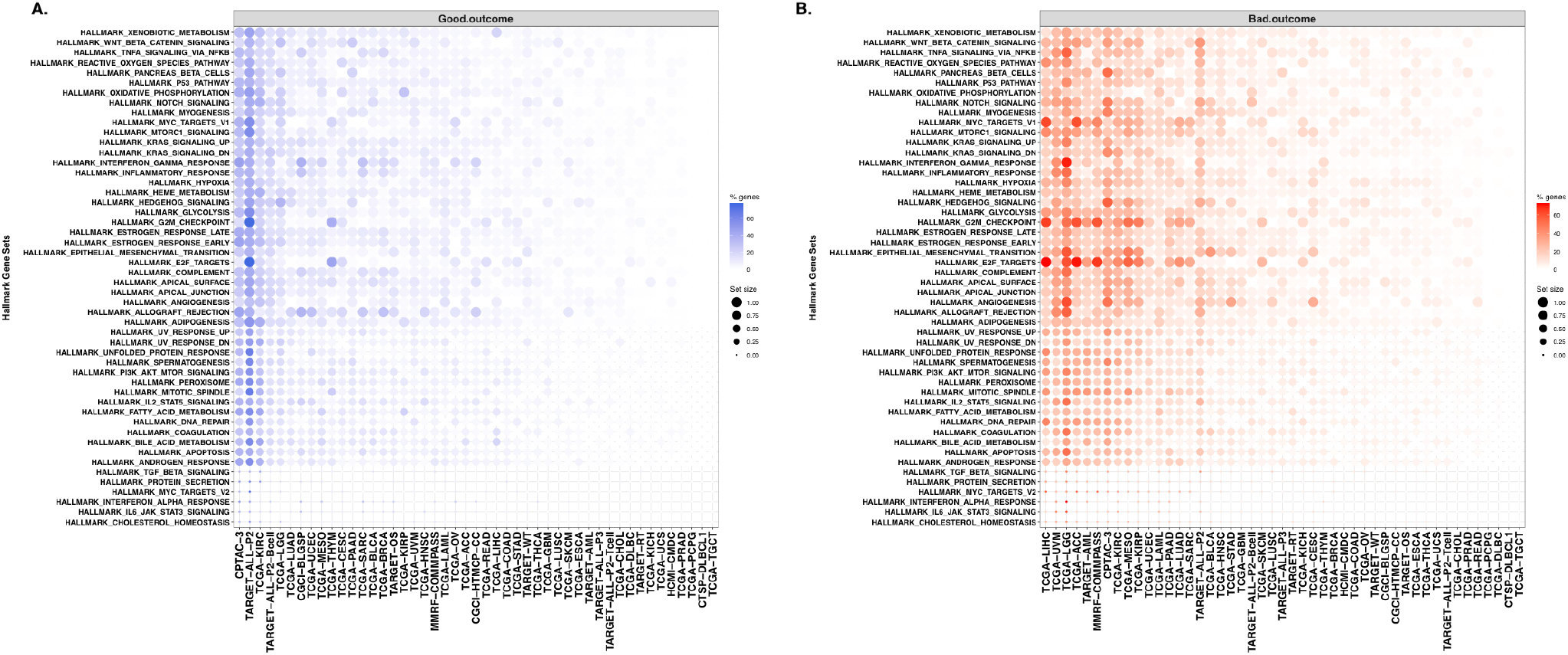
Hallmark genesets with enrichment of survival associated genes. Summary of significantly associated hallmark genes with good (A) and bad (B) overall survival for each hallmark gene set for each cancer type. The dots color gradient represents the percentage of genes per gene set (% genes) that are significantly associated (p<0.05) with survival outcome, while size of the dots represents the total number of genes within a gene set (set size). The blue and orange color gradient indicate the good (HR<1) and bad (HR>1) survival outcomes.

The top cancer types that contained the highest proportion of hallmark genes significantly associated with good overall survival are TARGET-ALL-P2, CPTAC-3, TCGA-KIRC **(Fig 5A)**. On the other hand, most of the hallmark genes depicted association with poor outcomes in TCGA-LGG, TCGA-LIHC, TCGA-UVM, TCGA-ACC cancers (Fig 5B). Regardless of the cancer types, genes such as *ITGA5, SERPINE1, EIF4EBP1* are majorly associated with bad outcomes; while genes such as *ALDH2, DBP, FRAT1, PDCD4* are associated with good outcomes.

Single-gene analysis of low-grade gliomas (TCGA-LGG) that showed poor prognosis at the gene set level depicted approximately 30-40% genes associated with poor outcomes (hazard Ratio >1 and P value <.05) (Fig 5B). These genes belong mainly to immune response and inflammatory gene sets such as HALLMARK inflammatory response, HALLMARK interferon-gamma response, HALLMARK TNFA signaling via NF-KB, and HALLMARK IL2 STAT5 signaling **(Fig 5B).**

Similarly single-gene analysis of TARGET pediatric acute lymphoid leukemia (TARGET-ALL-P2) showed that 40-50% from hallmark genesets correlated with good outcome **(Fig 5A)**. Majority of these genes belong to metabolic and inflammatory gene sets such as HALLMARK interferon gamma response, HALLMARK glycolysis, HALLMARK Kras signaling down, and HALLMARK adipogenesis.

#### Use Case 3: *Survival analysis of T cell exhaustion to effector marker genes ratio*

To determine the impact of the ratio of exhausted and functional T cells on cancer outcomes, we performed gene ratio-based analysis using the Survival Genie platform. For the analysis, we have used expression levels of Layilin (LAYN), a potent maker for quantifying tumor-infiltrating exhausted CD8^+^ T cells ^25^. We have chosen GZMA and IFNϒ as markers for quantifying functional and activated CD8+ T cells ^26^. We explored the effect of exhausted to effector CD8 T cell ratio on survival outcome across adult and pediatric cancers.

The RNA-seq normalized expression values were used to compute the ratios of LAYN to GZMA and LAYN to IFNG for each tumor sample. The results of cox regression analysis across 47 cancer types for each gene ratio, i.e., LAYN: GZMA and LAYN: IFNG are shown in **Fig. 6.** The higher LAYN: GZMA ratio **(Fig. 6A)** showed a significant association with poor survival in bladder, breast, head and neck, lung, liver, stomach, thyroid, and uterine cancer (Supplementary Table 4); while low LAYN: GZMA expression ratios were significantly associated with better prognosis in multiple myeloma, leukemia, low-grade glioma, and kidney cancer. A similar significant association of high LAYN to IFNG ratio with poor survival could also be observed in CPTAC-3, TCGA-BLCA, TCGA-BRCA, TCGA-HNSC, TCGA-UCEC **(Fig. 6B)**. On the contrary, in datasets from TCGA-KIRC, TCGA-KIRP, and TCGA-UVM higher ratios of LAYN: IFNϒ was associated with better survival. These data suggest that high expression of exhausted T cells marker (or lower expression of effector T cell marker) tend to be associated with a poor prognosis. However, high expression of cytolytic markers, GZMA or IFNϒ tend to suggest a good prognosis in blood and kidney cancers, respectively.

**Figure 6.**
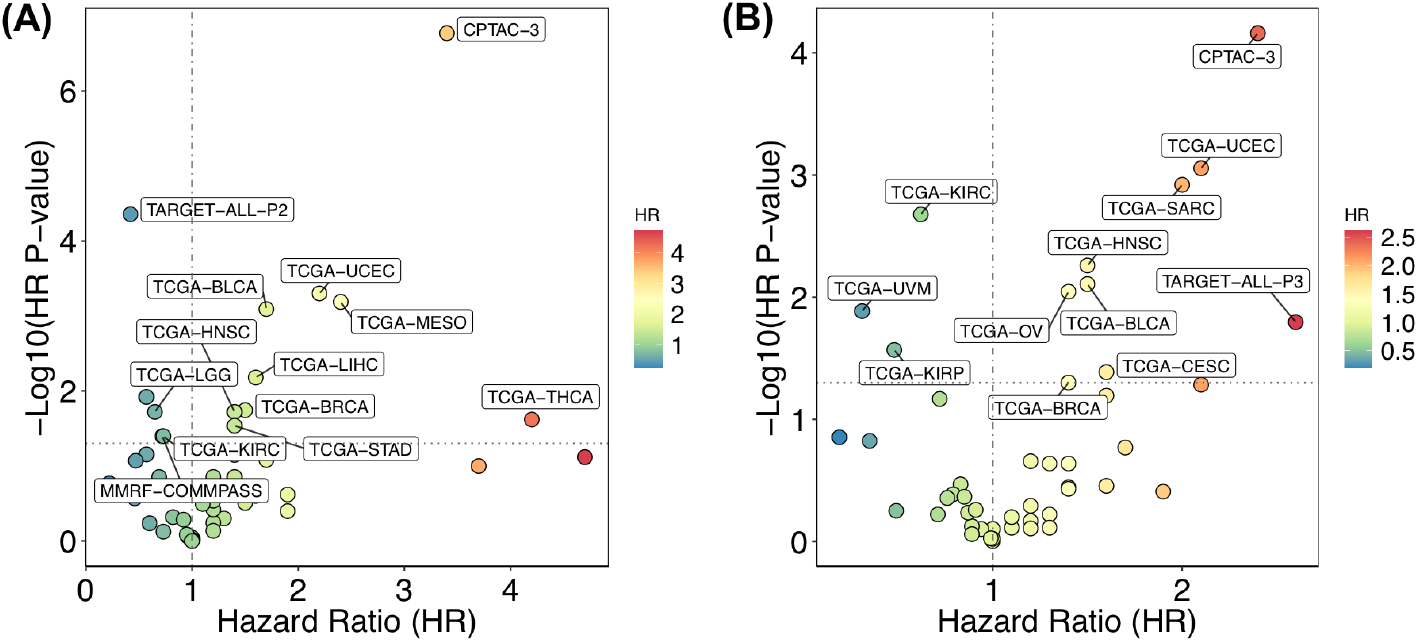
Gene Ratio based survival analysis. Association of (A) LAYN: GZMA and (B) LAYN: IFNG gene ratios on patient survival in different types of cancer. Only the significantly associated cancer types are labeled.

#### Use Case 4: *Survival analysis of single-cell RNA-seq T cell clusters*

We illustrate the utility of Survival Genie for scRNA-seq cluster markers prognostic utility by implementing it for survival analysis on T cell clusters **(Fig.7A)** generated from scRNA-seq analysis of paired pediatric AML bone marrow samples taken at the time of diagnosis and end of induction ^27^. We performed Cox proportional survival analysis using TCGA-LAML dataset **(Fig. 7).** Out of the 11 T cell clusters, the relapse-associated cluster (#CL11) depicted significant association with poor survival (HR=2, p=0.001) **(Fig 7B)**. Moreover, cluster 11 (#CL11) depicted overexpression of CD69, a type II glycoprotein that is known to regulate inflammation and exhaustion of tissue-resident T cells and promote tumor growth/relapse ^28^.

**Figure 7.**
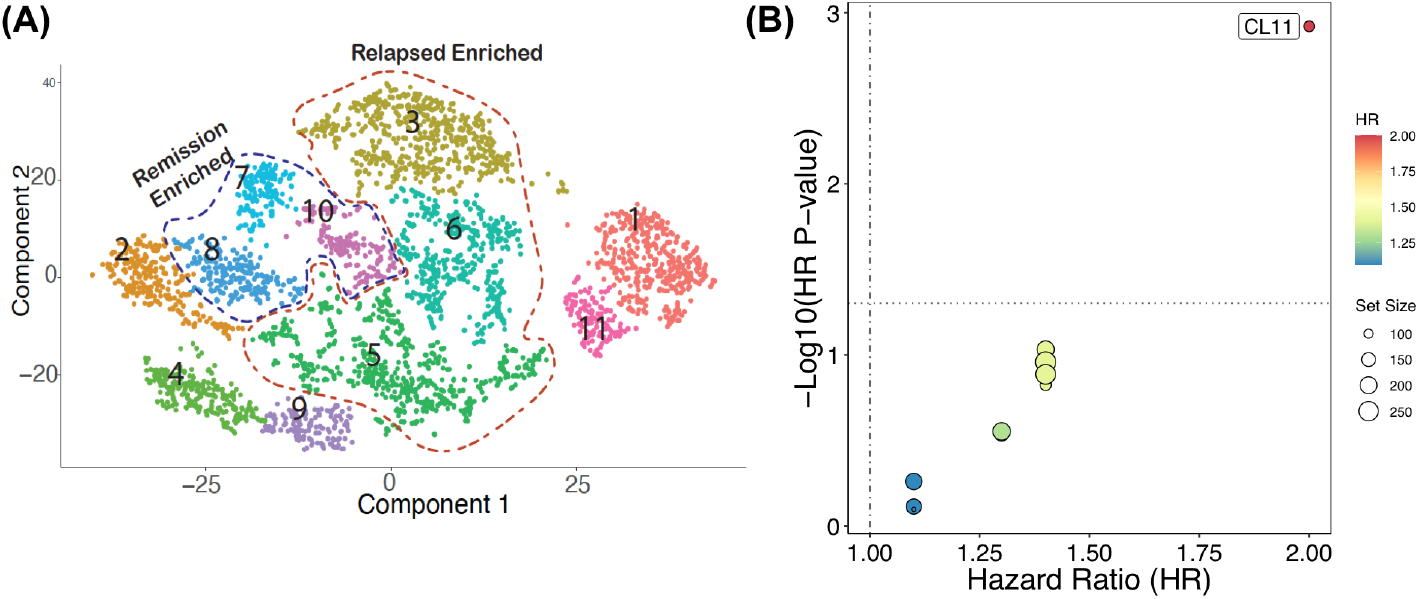
Single-cell clusters based survival analysis. (A) UMAP of single-cell RNA-seq data from relapse-associated non-AML cells at the time of diagnosis. The colors represent the 11 distinct sub-clusters of T cells. (B) Prognostic significance of identified scRNA-seq clusters on overall survival in TCGA-LAML patient dataset. The significant associations between cluster enrichment and OS are highlighted.

## DISCUSSION

In this genomics era with abundant cancer genomics data, the bioinformatics tools to explore the prognostic utility of individual genes or gene sets are a major bottleneck in identifying robust prognostic biomarkers using the power of big data. Despite the multiple existing survival analysis tools, these tools have multiple shortcomings including data comprehensiveness and survival analysis on gene markers from single-cell data or cell fractions. Using traditional transcriptomic data, a plethora of individual genes have been tested and numerous suggested as potential prognosis markers for different cancer. As intuitively obvious to calculate single sample enrichment scores for a gene set per sample to test prognostic utility, it is not routinely used for survival analysis. With Survival Genie, researchers can easily explore multi-gene markers at once and identify gene sets associated with cancer survival. The analytical options in Survival Genie makes it a unique resource for predicting clinical response based on profiles other than single-gene or average multiple-genes expression across different cancers with extended functionality to evaluate survival associations of single-cell clusters.

With multiple data sources and no consensus on cross-platform data processing and normalization, data standardization and optimization, cohort size, and analysis workflows, different survival tools yield different results. To maintain uniformity in the genomic data, while developing Survival Genie we have used harmonized cancer datasets from the NCI Genomic Data Commons. At present, GDC^4^ contains multidimensional datasets from 68 cancer programs, all processed and analyzed with latest human reference genome build GRCh38 and up-to-date workflows. This uniformly processed data allows comparative analysis of prognostic genes or genesets with minimal bias resulting from inconsistent data preprocessing, normalization, or technological differences among datasets. To further enhance the comprehensiveness of Survival Genie, we are planning to include data from other cancer genomics initiatives including cancer projects such as BEATAML1.0^29^, Cancer Genome Characterization Initiative (CGCI), and Cancer Dream Team projects.

Exploring the prognostic utility of different malignant and immune cells clusters from single-cell data allows exploration of prognostic utility of heterogeneous malignant tumor cells along with the immune microenvironment. In the bulk RNA-seq data we may not be able to determine how single-cell signatures are related to cell level composition during tumor progression. This certainly holds true when studying rare sub clonal profiles in cancers such as acute myeloid leukemia. To address this limitation, we estimated the cellular composition of tumors from bulk RNA-seq expression by deconvolution, using immune cell-type specific profiles from tumor-derived single-cell RNA sequencing data^21^. The resulting cell correlation matrix can subsequently be used to identify the cell type abundance in bulk tissue data when predicting outcomes.

We have improved upon the existing tools in the ability to predict prognosis from multi-gene marker sets. In the future, efforts will be made to include options for multivariable regression and in-depth subtype analysis using clinicopathological features (tumor grade, lymph node status, treatment, mutation status, subtypes, tumor tissue image, microsatellite instability). In addition, Survival Genie will also allow exploration of prognostic associations of miRNA, long non-coding RNA and hyper/hypo methylated genomics regions. We also intend to include the ability to subset survival analysis based on genes that correlate with canonical cell specific markers (e.g., T-cells), thus allowing survival analysis based on cell-specific signatures. Survival Genie is an open-source webbased tool running and being tested since September 2020 and we intend to maintain it for a minimum of five years.

## CONCLUSION

In summary, Survival Genie allows the exploration of prognostic potential of genes and gene sets in a broad range of cancer datasets and specifically to validate the cell-type specific markers from single-cell mRNA sequencing. The exploration of three-way relationship between the cell-specific biomarker genes, enrichment of cell types and survival outcomes will assist in understanding how specific gene impacts cellular composition tumor microenvironment to drive cancer outcomes. This will assist in developing biomarkers to predict the immune-suppressive microenvironment that might lead to identification of next-generation candidates for targeted immunotherapies.

## ACKNOWLEDGEMENTS

We would like to thank the Winship Research Informatics shared resource of the Winship Cancer Institute of Emory University for supporting the web server. Thanks to Beena E. Thomas, and Gulay B. Ulukaya for reviewing and editing the manuscript.

## DECLARATIONS

### Ethical Approval

None

## Supplementary Tables Information

**Supplementary Table 1.** List of the cancer datasets along with their data types available in the Survival Genie.

**Supplementary Table 2:** Survival Associations of hallmark gene sets (n=50) and cancers.

**Supplementary Table 3:** Results of survival analysis across 47 cancer datasets for individual gene from hallmark gene set.

**Supplementary Table 4:** Ratio based survival analysis of T cell exhaustion to effector marker genes across different cancer datasets.

## REFERENCES

1 Cancer Genome Atlas Research, N. et al. The Cancer Genome Atlas Pan-Cancer analysis project. Nat Genet 45, 1113–1120, doi:10.1038/ng.2764 (2013).

2 International Cancer Genome, C. et al. International network of cancer genome projects. Nature 464, 993–998, doi:10.1038/nature08987 (2010).

3 Ma, X. et al. Pan-cancer genome and transcriptome analyses of 1,699 paediatric leukaemias and solid tumours. Nature 555, 371–376, doi:10.1038/nature25795 (2018).

4 Grossman, R. L. et al. Toward a Shared Vision for Cancer Genomic Data. N Engl J Med 375, 1109–1112, doi:10.1056/NEJMp1607591 (2016).

5 Zheng, H. et al. Comprehensive Review of Web Servers and Bioinformatics Tools for Cancer Prognosis Analysis. Front Oncol 10, 68, doi:10.3389/fonc.2020.00068 (2020).

6 Cerami, E. et al. The cBio cancer genomics portal: an open platform for exploring multidimensional cancer genomics data. Cancer Discov 2, 401–404, doi:10.1158/2159-8290.CD-12-0095 (2012).

7 Chandrashekar, D. S. et al. UALCAN: A Portal for Facilitating Tumor Subgroup Gene Expression and Survival Analyses. Neoplasia 19, 649–658, doi:10.1016/j.neo.2017.05.002 (2017).

8 Borcherding, N., Bormann, N. L., Voigt, A. P. & Zhang, W. TRGAted: A web tool for survival analysis using protein data in the Cancer Genome Atlas. F1000Res 7, 1235, doi:10.12688/f1000research.15789.2 (2018).

9 B., G. Survival analysis across the entire transcriptome identifies biomarkers with the highest prognostic power in breast cancer. Computational and Structural Biotechnology Journal 19, 4101–4109 (2021).

10 Gyorffy, B. et al. An online survival analysis tool to rapidly assess the effect of 22,277 genes on breast cancer prognosis using microarray data of 1,809 patients. Breast Cancer Res Treat 123, 725–731, doi:10.1007/s10549-009-0674-9 (2010).

11 Wang, F. et al. OSuvm: An interactive online consensus survival tool for uveal melanoma prognosis analysis. Mol Carcinog 59, 56–61, doi:10.1002/mc.23128 (2020).

12 Pak, K. et al. A User-Friendly, Web-Based Integrative Tool (ESurv) for Survival Analysis: Development and Validation Study. J Med Internet Res 22, e16084, doi:10.2196/16084 (2020).

13 GenomicDataCommons: NIH / NCI Genomic Data Commons Access (2021).

14 Clark, K. et al. The Cancer Imaging Archive (TCIA): maintaining and operating a public information repository. J Digit Imaging 26, 1045–1057, doi:10.1007/s10278-013-9622-7 (2013).

15 A Package for Survival Analysis in R (2021).

16 Bioconductor Open Source Software for Bioinformatics v. 3.13.

17 R: A language and environment for statistical computing. (R Foundation for Statistical Computing, 2017).

18 Shiny R web-server.

19 Barbie, D. A. et al. Systematic RNA interference reveals that oncogenic KRAS-driven cancers require TBK1. Nature 462, 108–112, doi:10.1038/nature08460 (2009).

20 Hao, Y. et al. Integrated analysis of multimodal single-cell data. Cell 184, 3573–3587 e3529, doi:10.1016/j.cell.2021.04.048 (2021).

21 Chen, B., Khodadoust, M. S., Liu, C. L., Newman, A. M. & Alizadeh, A. A. Profiling Tumor Infiltrating Immune Cells with CIBERSORT. Methods Mol Biol 1711, 243–259, doi:10.1007/978-1-4939-7493-1_12 (2018).

22 survMisc: Miscellaneous Functions for Survival Data (2018).

23 Liberzon, A. et al. The Molecular Signatures Database (MSigDB) hallmark gene set collection. Cell Syst 1, 417–425, doi:10.1016/j.cels.2015.12.004 (2015).

24 Liberzon, A. et al. Molecular signatures database (MSigDB) 3.0. Bioinformatics 27, 1739–1740, doi:10.1093/bioinformatics/btr260 (2011).

25 Zheng, C. et al. Landscape of Infiltrating T Cells in Liver Cancer Revealed by Single-Cell Sequencing. Cell 169, 1342–1356 e1316, doi:10.1016/j.cell.2017.05.035 (2017).

26 Roufas, C. et al. The Expression and Prognostic Impact of Immune Cytolytic Activity-Related Markers in Human Malignancies: A Comprehensive Metaanalysis. Front Oncol 8, 27, doi:10.3389/fonc.2018.00027 (2018).

27 Thomas, B. P. P.; Bhasin, S.; Sarkar, D.; Dwivedi, B.; Park, S.; DeRyckere, D.; Henry, C.; Raikar, S.; Porter, C.; Pauly, M.; Summers, R.; Castellino, S.; Wechsler, D.; Graham, D.; Bhasin, M.. Single Cell Transcriptomics Revealed AML and Non-AML Cell Clusters Relevant to Relapse and Remission in Pediatric AML. Blood 136(Supplement 1), 24–25, doi:10.1182/blood-2020-142513 (2020).

28 Mita, Y. et al. Crucial role of CD69 in anti-tumor immunity through regulating the exhaustion of tumor-infiltrating T cells. Int Immunol 30, 559–567, doi:10.1093/intimm/dxy050 (2018).

29 Tyner, J. W. et al. Functional genomic landscape of acute myeloid leukaemia. Nature 562, 526–531, doi:10.1038/s41586-018-0623-z (2018).

